# Host-dependent differences in replication strategy of the *Sulfolobus* Spindle-shaped Virus strain SSV9 (a.k.a., SSVK1): Lytic replication in hosts of the family Sulfolobaceae

**DOI:** 10.1101/2020.03.30.017236

**Authors:** Ruben Michael Ceballos, Coyne Drummond, Carson Len Stacy, Elizabeth Padilla Crespo, Kenneth Stedman

## Abstract

The *Sulfolobus* Spindle-shaped Virus (SSV) system has become a model for studying thermophilic virus biology, including archaeal host-virus interactions and biogeography. Several factors make the SSV system amenable to studying archaeal genetic mechanisms (e.g., CRISPRs) as well as virus-host interactions in high temperature acidic environments. First, it has been shown that endemic populations of *Sulfolobus*, the reported SSV host, exhibit biogeographic structure. Second, the acidic (pH<4.5) high temperature (65-88°C) SSV habitats have low biodiversity, thus, diminishing opportunities for *host switching*. Third, SSVs and their hosts are readily cultured in liquid media and on gellan gum plates. Fourth, given the wide geographic separation between the various *SSV-Sulfolobus* habitats, the system is amenable for studying allopatric versus sympatric virus-host interactions. Previously, we reported that SSVs exhibit differential infectivity on allopatric and sympatric hosts. We also noticed a wide host range for virus strain SSV9 (a.k.a., SSVK1). For decades, SSVs have been described as “non-lytic” dsDNA viruses that infect species of the genus *Sulfolobus* and release virions via “blebbing” or “budding” as a preferred strategy over host lysis. Here, we show that SSVs infect more than one genus of the family Sulfolobaceae and, in allopatric hosts, SSV9 does not appear to release virions by blebbing. Instead, SSV9 appears to lyse all susceptible allopatric hosts tested, while exhibiting canonical non-lytic viral release via “blebbing” (historically reported for all other SSVs), on a single sympatric host. Lytic versus non-lytic virion release does not appear to be driven by multiplicity of infection (MOI). Greater relative stability of SSV9 compared to other SSVs (i.e., SSV1) in high temperature, low pH environments may contribute to higher transmission rates. However, neither higher transmission rate nor relative virulence in SSV9 infection drives replication profile (i.e., lytic versus non-lytic) in susceptible hosts. Although it is known that CRISPR-Cas systems offer protection against viral infection in prokaryotes, CRISPRS are not reported to be a determinant virus replication strategy. Thus, the genetic/molecular mechanisms underlying SSV9-induced lysis are unknown. These results suggest that there are unknown genetic elements, resulting from allopatric evolution, that drive virion release strategy in specific host strain-SSV strain pairings.

## INTRODUCTION

Reduced model systems have been used extensively to study fundamental properties of virus evolution (Lenski and Levin, 1985; Morgan et al., 2005; Brockhurst et al., 2007). Moreover, it suggested that studying virus-host interactions in reduced model systems may provide opportunities to understand fundamental processes of virus evolution in host systems of agricultural or medical importance (Brockhurst et al., 2007; Dennehy, 2009). Given the recent view that the evolutionary origin(s) of viruses may be linked to the early evolution of Archaea (Forterre et al., 2006; Berliner et al., 2018) and that the emergence of viruses (not bacteriophage) likely pre-dates the divergence of the Archaea and Eukarya (Krupovic et al., 2017; Prangishvili et al., 2017), a robust archaeal virus model system could provide new insights into the evolution of virus lineages and viral replication strategies as well as mechanisms of viral virulence, host resistance, and virus attenuation.

The *Sulfolobus* Spindle-shaped Virus (SSV) system has become a popular model for studying thermophilic archaeal virus biology and virus-host biogeography. Several factors make this system ideal for studying virus-host infections in crenarchaea (i.e., Sulfolobales). First, endemic populations of SSV hosts from the family Sulfolobaceae exhibit biogeographic structure such that there is a positive correlation between genetic distance among strains (i.e., divergence) and geographic distance between the sites from which strains have been isolated (Grogan et al., 1989; Whitaker et al., 2003; Reno et al., 2009). SSVs also exhibit biogeographic structure on a global-scale (Held and Whitaker, 2009). Second, the highly acidic (pH<4.5) and high temperature (65°C-88°C) SSV-*Sulfolobus* habitats feature low biodiversity, limiting the potential for host switching, which can confound efforts to elucidate the genetic underpinnings of virus-host infection profiles (Munson-McGee et al., 2018). Third, SSVs and Sulfolobales can be readily cultured both in liquid media (e.g., yeast-sucrose, tryptone) and on gellan gum (e.g., Gel-Rite^®^) plates (Zillig et al., 1996; Stedman, 2008; Ceballos et al., 2012). Fourth, given the wide geographical separation between the sulfuric geothermal areas, which are habitats for SSVs and their hosts, this system (Fig. 1) is amenable to studying multiple allopatric and sympatric virus-host pairs (Ceballos et al., 2012), an essential for studying virus-host interactions, biogeography, and coevolution (Greischar & Koskella, 2007).

**FIGURE 1.**
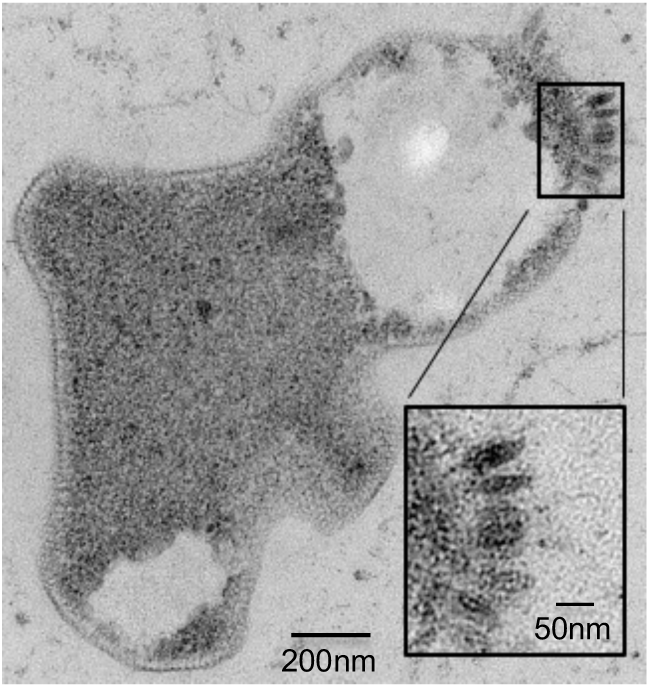
SSV infection of a Sulfolobus host. Transmission Electron Micrograph (TEM) of a 30-60μm epoxy-impregnated section/slice from a SSV-infected cell. (60,000X magnification). Vacuole-like anomalies in the intracellular space and membrane disruptions are observed in SSV infected host cells. A closer view (inset) reveals fusiform or “spinde-shaped” virus-like particles on the surface of the cell membrane.

For over three decades, SSVs were considered to be non-lytic ‘*blebbing’* or budding viruses that exclusively infect *Sulfolobus* (Martin et al., 1984; Reiter et al., 1987; Palm et al., 1991; Schleper et al., 1992; Zillig et al., 1998; Wiedenheft et al., 2004; Contursi et al., 2006; Prangishvili et al., 2006; Ceballos et al., 2012; Fusco et al., 2015; Quemin et al., 2016). The term *non-lytic* refers to the fact that SSV infection results in inhibition of cell growth (both in liquid culture and on lawns) rather than the gross lysis of host cells and cell death in liquid culture or clear plaques on host lawns - both of which result from lytic replication. Recently, it was suggested that SSV9 can induce a state of dormancy, empty cells, and eventual host death in a sympatric host; however, the mechanism for this proposed dormancy is unclear and there does not seem to be any cell lysis (Bautista et al., 2015).

Plaque-like “halo” assays using both sympatric and allopatric hosts have repeatedly demonstrated that different SSVs (e.g., SSV1, SSV2, SSV3, SSV8, SSV10) form turbid indentations or *halos* on host lawns (Fig. 2A, B), which often feature diffuse boundaries (Martin et al., 1984; Schleper et al., 1992; Wiedenheft et al., 2004; Ceballos et al., 2012; Iverson and Stedman, 2012). Yet, halo assays using SSV9 (formerly known as SSVK1), isolated from the Valley of Geysers (*Russian*: Дoлина гейзеров) Kamchatka, Russia (Wiedenheft et al., 2004), form large *clear* plaques on host lawns (*see* Fig. 2C) in contrast to the *turbid* diffusely-bound halos characteristic of all other SSVs (Ceballos et al., 2012), indicating that SSV9 may replicate differently than the other SSVs (Bautista et al., 2015).

**FIGURE 2.**
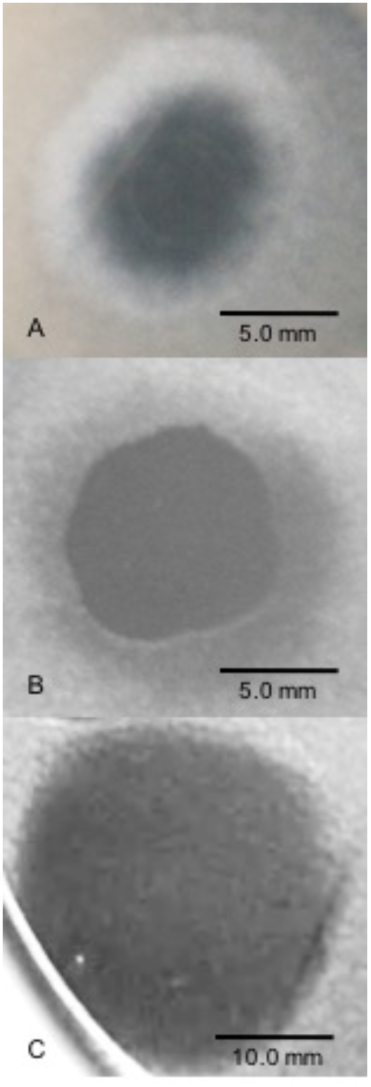
SSV Spot-On-Lawn Halo Assays. A 1-2μL drop of virus suspension was spotted on lawns of DSM 1617 (formerly, *Sulfolobus solfataricus* strain P2) to determine whether a viral infection can be established and the nature of the virus-host interaction in terms of halo “phenotype”. (A) SSV1 forms broad diffuse, turbid halo. (B) SSV8 (a.k.a., SSVRH) makes a broad diffuse, turbid halo with a sharper boundary between the direct application point and halo leading edge; (C) SSV9 (a.k.a., SSVK1) yields a large and complete clearing of host lawn.

To test the hypothesis: SSV9 lyses susceptible hosts of the family *Sulfolobaceae* - host growth in single-virus/single-host interactions using three distinct SSV strains (i.e., SSV1, SSV8, and SSV9) was evaluated in both liquid culture and with *spot-on-lawn “*halo” assays. Both allopatric and sympatric hosts of the family *Sulfolobaceae* (Order: Sulfolobales) were used to determine relative susceptibility. Several different liquid culture conditions were used, specifically: a variety of distinct virus strain-host strain pairings were examined; different multiplicities of infection (MOI) were tested; and, end-point assays measuring virus and host dynamics in parallel were conducted. Spot-on-lawn halo assays and observations from small-scale liquid culture infections were conducted to examine plaque-like halo formation and the relative amount of cell debris emerging from different virus-host strain pairings, respectively. The relative stability of different SSV strains under conditions that simulate the natural environment of these viruses and their hosts were also conducted to assess whether virion stability plays a role in successful transmission (and thus between strain differences in virion production).

## RESULTS

### Virus Yields and Confirmation of SSV Infection

Titers of virus stocks were determined by one (or more) of four distinct methods: electrospray ionization/mass spectrometry (ESI/MS) in units of virus particles per milliliter (VP/mL); serial dilution plaque-like assays in units of halo-forming units per milliliter (hfu/mL); virus-like particle (VLP) counts using transmission electron micrographs (VLPs/mL); and/or, quantitative polymerase chain reaction (qPCR) in terms of the total number of viral genomes per milliliter (v_g_/mL). For infection assays, end-point titers were also determined by one or more of these methods with ESI/MS and plate assays preferred.

Growing 600mL liquid cultures of permissive host, each infected with a single SSV strain (i.e., SSV1, SSV8, or SSV9), to an optical density (OD_600nm_) of 0.8 (∼5.4×10^8^ cells/mL) results in comparable virion yields as shown by electrospray ionization/mass spectrometry (ESI/MS) spectra (Fig. S1). Upon: harvest, centrifugation, removal of supernatant, supernatant filtration (0.45μm) and concentration by ultrafiltration (∼300-fold) - SSV8 and SSV9 suspensions result in ∼2×10^8^ – 4×10^8^ VP/mL. SSV1 yields are typically 50-80% lower. Thus, it takes 1.2-2.0L of SSV1-infected culture to ensure stocks are of the same order of magnitude and comparable to SSV8 and SSV9. SSV suspensions were then diluted to produce working stocks with equivalent titers (e.g., ∼1×10^8^ VP/ml). These working stocks were used for comparative infection assays with the same multiplicity of infection (MOI). MOIs of 0.01 and 0.001 were typically used but other MOIs (e.g., MOI=3) were also tested.

It has previously been reported that not all species within the family *Sulfolobaceae* are susceptible to SSV infection (Ceballos et al., 2012). Thus, to ensure that there was active infection in the virus-host pairings selected for this study, three steps were taken. First, virus suspensions were examined under transmission electron microscopy (TEM) to ensure that spindle-shaped virus particles (i.e., virions) were present in the harvested virus stocks. Second, *spot-on-lawn* plate assays were used as a qualitative (i.e., yes or no) verification that a given SSV strain is able to infect a specific strain within the family *Sulfolobaceae*. Third, post-infection titers and TEM were performed to verify active infection (Fig. 3).

**FIGURE 3.**
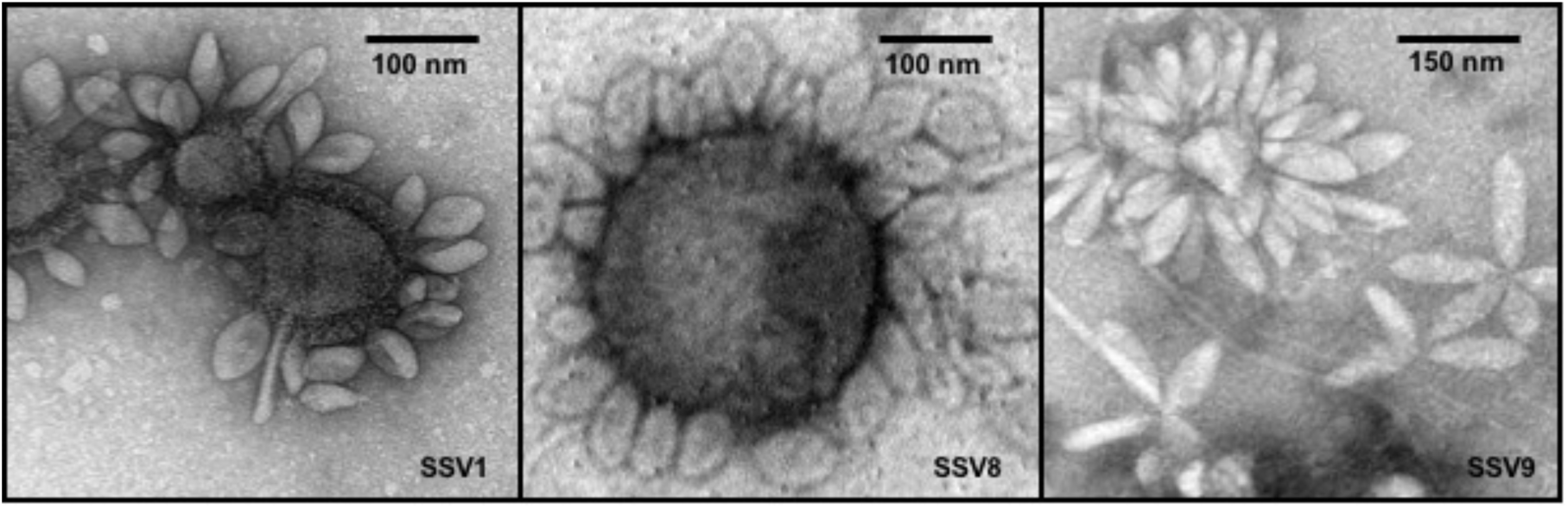
Transmission Electron Micrographs of SSV Particles. Transmission Electron Microscopy (TEM) demonstrates the presence of spindle-shaped virus particles after harvest and concentration steps at end-points of infection assays as well as during the preparation of virus stocks for liquid culture infection assays and halo assays. TEM images of SSV1 [*left*], SSV8 [*middle*], and SSV9 [*right*].

### Host-Growth after Infection by Various SSVs

Liquid culture assays using several SSVs infecting the universally susceptible host strain *Sulfolobus* strain Gθ (Cannio et al., 1998; Ceballos et al., 2012), reveal that host growth (μ_max_) is slowed and the “carrying capacity” upon entering stationary phase (i.e., *N*_*asymptote*_) is reduced compared to uninfected control (Fig. 4). Individual (i.e., single virus-single host) infections with SSV1, SSV8, or SSV10 on strain Gθ show archetypal host growth with these reportedly non-lytic SSVs. Infection with either of two SSVs isolated from two geographically-distinct Icelandic geothermal regions - SSV2 (from Reykjanes) and SSV3 (from Krisovic) - also show reductions in μ_max_ and area-under-the curve (AUC) when compared to controls. Despite SSV2 and SSV3 infections exhibiting slightly different host growth profiles (Fig. 4A, cyan and mustard lines) due to slower growth to stationary phase, data still fit to a Gompertz model with high coefficient of determination values (Table S1). Measures of percent inhibition can be derived by area-under-the-curve (AUC) calculations for the infected host growth and dividing by the AUC for the uninfected control (Fig. 4B). Using this quotient, an index of relative virulence (*V*_*R*_) between strains on a given host can be generated (Stacy and Ceballos, 2020).

**FIGURE 4.**
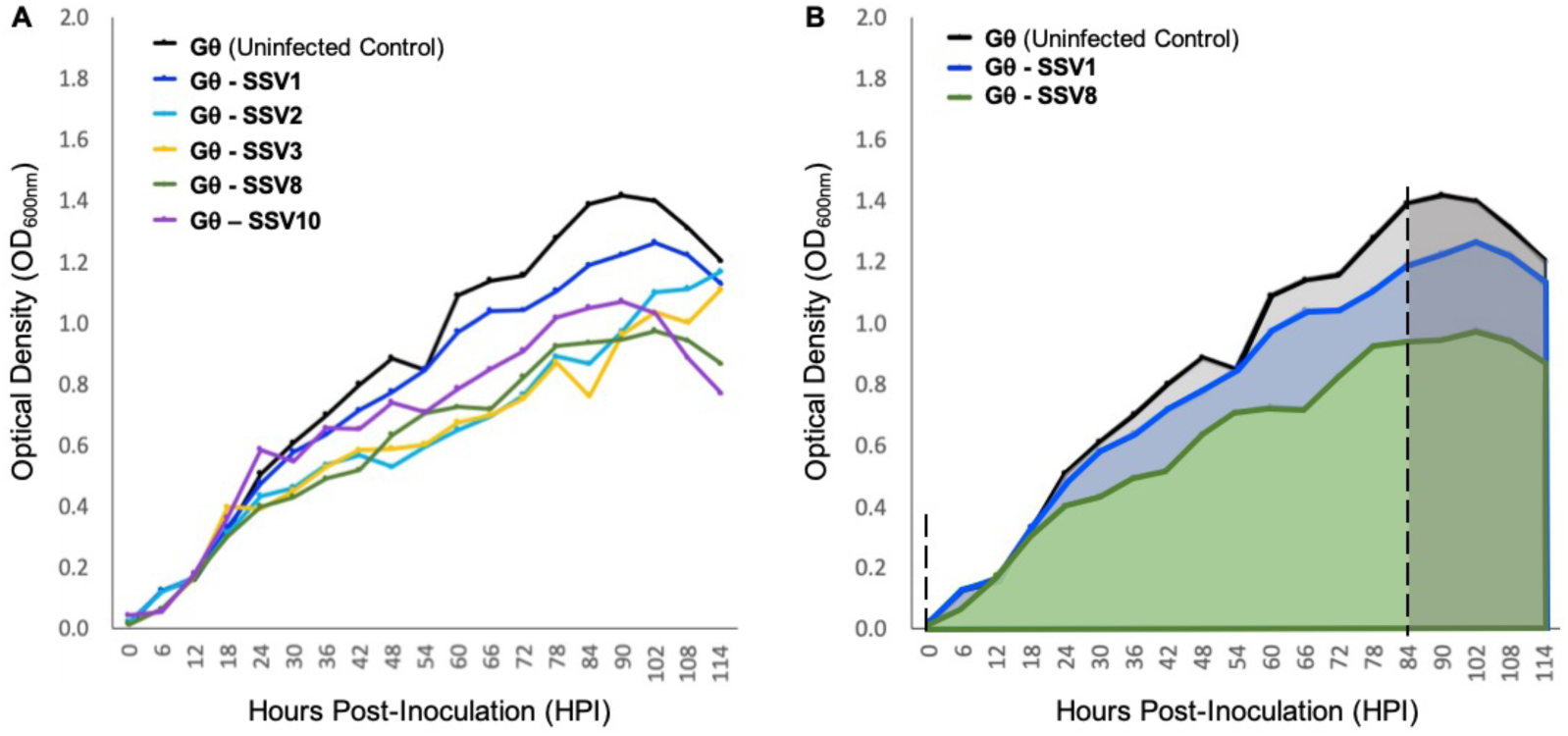
Growth of host strain *Sulfolobus* Gθ during SSV infection. (A) *Sulfolobus* strain Gθ challenged with SSV1 (Palm et al., 1991), SSV2 (Stedman et al., 2003), SSV3 (Stedman et al., 2006), SSV8 (a.k.a., SSVRH; Wiedenheft et al., 2004) and SSV10 (a.k.a., SSVL1; Goodman & Stedman, 2018). (B) Graphical representation of percent inhibition of SSV1 and SSV8 using area-under-the-curve (AUC). All growth curves fit with high coefficients of determination (R2 values) to Gompertz Models or Logistic Growth Models (*see* Table S1). The asymptotes (dashed) indicate point of inoculation until end of the exponential growth phase/onset of stationary phase. Infections were conducted at an MOI = 0.01.

Using a Gompertz model and AUC as a measure of percent inhibition (PI), SSV1 exhibits the lowest PI (8.57%) while SSV8 has a significantly higher PI (28.44%) with R^2^ values for the Gompertz of 0.9875 and 0.9775, respectively. Employing Riemann Sums or alternative models (i.e., Logistic Growth) does not significantly alter relative virulence (*V*_*R*_) for SSV1 and SSV8 (*see* Table S1). Regardless of analytical approach, the general trend in virulence of SSVs on Gθ is: SSV1<SSV10<<SSV2≈SSV3<SSV8. Note that AUC measures would suggest that strains SSV2 and SSV3 have a slightly greater PI than SSV8. However, note that AUC for SSV2 and SSV3 is underestimated due to a lag in both strains reaching stationary phase compared to the other SSVs (Stacy and Ceballos, 2020).

### Host-Growth after Infection with SSV9

Unlike infection with SSV1, SSV2, SSV3, SSV8 or SSV10, infecting *Sulfolobus* strain Gθ with SSV9 did not grow with Gompertzian-like (Gompertz, 1825; Laird, 1965) dynamics. In contrast, rapid and significant inhibition of culture growth was observed in cultures infected with SSV9. This pattern was observed for several strains of Sulfolobales over multiple trials with each strain (Fig. 5). Host strains tested were isolated from geographically-distinct geothermal regions from around the world and serve as allopatric hosts to SSV9.

**FIGURE 5.**
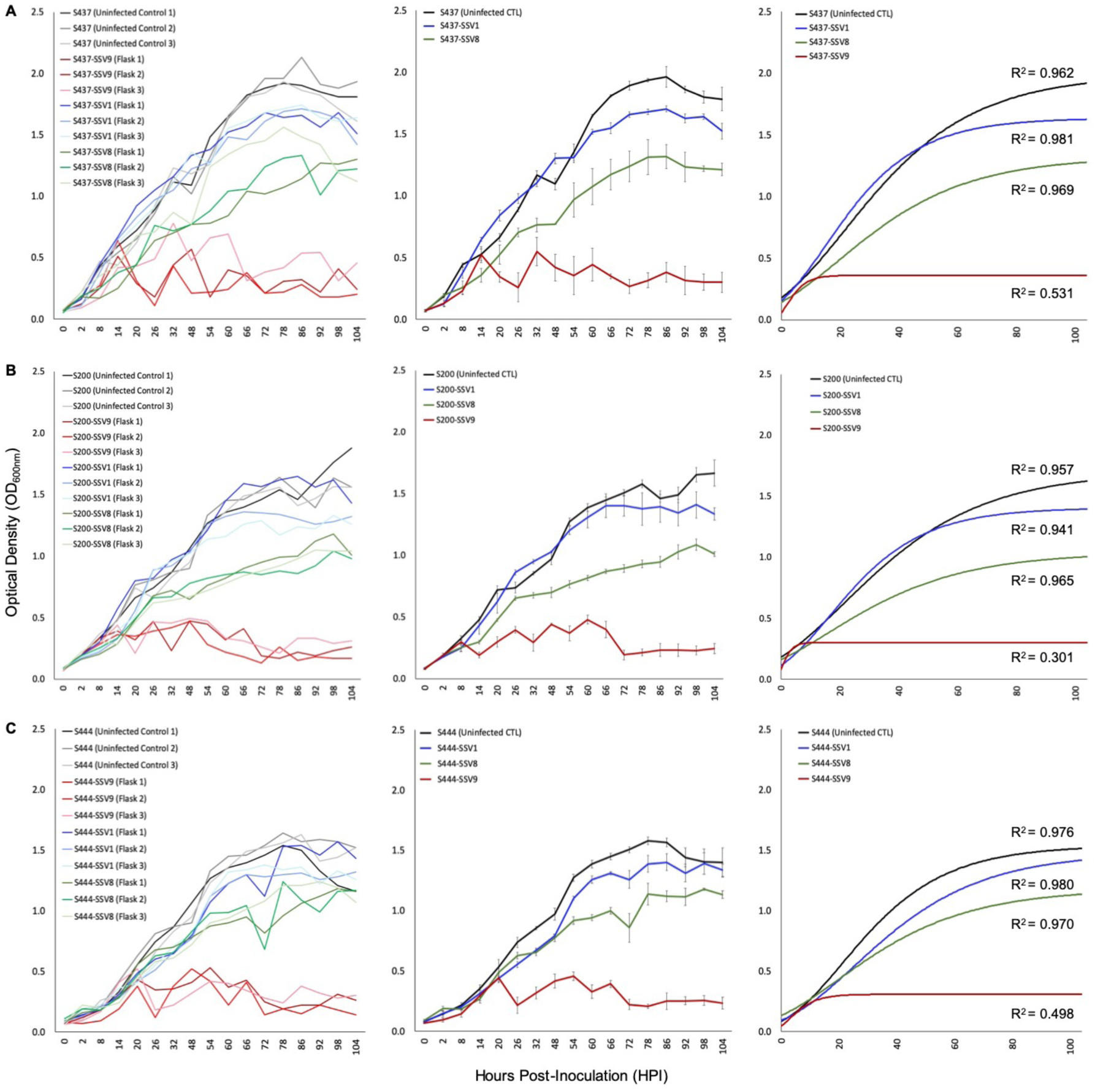
Growth Profiles for Susceptible Allopatric Host Strains Infected with SSVs. Under equivalent Multiplicity of Infection (MOI=0.1), allopatric hosts show classic Gompertzian growth when infected with either SSV1 or SSV8. However, cultures of the same hosts infected with SSV9 exhibit drastic growth inhibition. Growth in SSV9-infected cultures are “saw-toothed” with damped oscillations.

Liquid culture assays using SSV1, SSV8, and SSV9 on susceptible allopatric host strains: S437 (DSM1617 from the Deutsche Sammlung von Mikroorganismen und Zellculturen), a type strain (a.k.a., *S. solfataricus* P2) isolated from a hot spring near Pisciarelli, Italy; S200 (a.k.a., *S. icelandicus* HVE 10/4 from the Hveragerdi thermal region of Iceland); and, S444 (an isolate derived from Lassen Volcanic National Park, California, USA) – show reduced *N*_*asymp*_, AUC, and/or μ_max_ for SSV1 and SSV8 (versus uninfected controls). However, when each host was infected with SSV9, a distinct host growth profile emerged. Specifically, Gompertzian Models failed to adequately represent the SSV9 infection data. Instead, non-Gompertzian cyclical spike patterns are consistently observed in host growth. Low R^2^ values (<0.5) emerged for both individual traces and average curves for SSV9, whereas, high R^2^ values (> 0.9) were typical for all other SSVs infecting these same hosts (Table S2).

### Host-Growth Profile of SSV9 is not MOI-Dependent

To determine whether the unique growth profile of SSV9-infected hosts was dependent on MOI, dilutions of SSV8 and SSV9 stocks were prepared and liquid culture infection assays were performed on *Sulfolobus* strain Gθ. The typical host growth pattern shown for SSV infection (i.e., Gompertzian-like growth) is observed for all trials of SSV8 infection on strain Gθ regardless of MOI used (Fig. 6A). A positive correlation between SSV8 MOI and percent inhibition of host growth was generally observed. However, for the lowest MOI (1:64 and 1:256 dilutions), the PI was not significantly different from uninfected control despite productive infection. The atypical host growth profile for host infected with SSV9 also persisted (Fig. 6B). Amplitudes of host growth spikes change with an inverse relationship in response to MOI; however, there was no switch to the Gompertzian-like effect even at the lowest SSV9 MOI.

**FIGURE 6.**
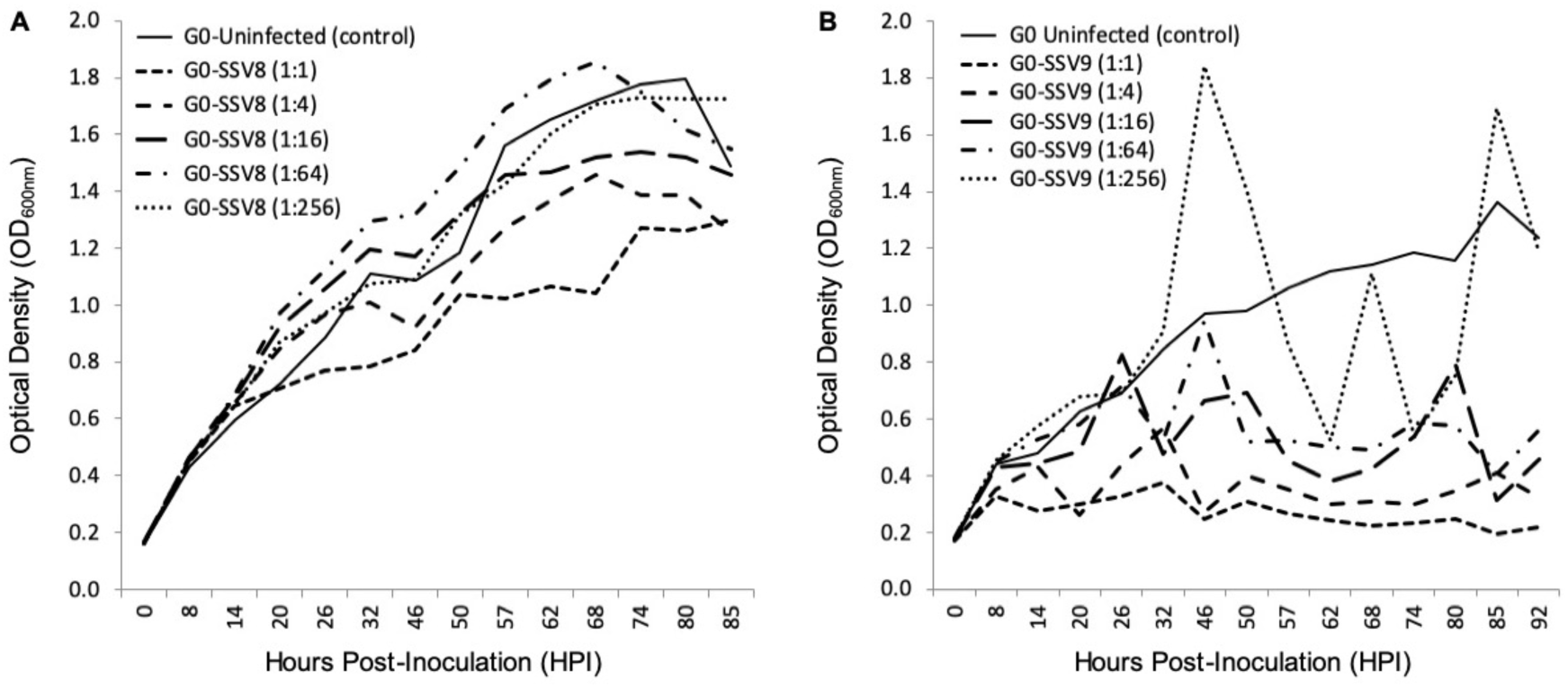
Growth of *Sulfolobus* strain Gθ Infected at Different MOIs: SSV8 versus SSV9. Under different Multiplicity of Infections, host strain Gθ (a.k.a., S13) infected with SSV9 retains the cyclical “saw-toothed” profile characteristic of SSV9 infection (and typical of lytic virus systems). Under different MOIs (1:1 represents a MOI = 0.1), strain Gθ infected with SSV8 retains Gompertzian growth.

### End-Point Infection Assays Indicate SSV9 Lytic Replication

To examine both host and virus population dynamics in a liquid culture infection assay, two large-scale trials (i.e., 14^+^ replicates) at higher observation frequency (every 4 hrs) were conducted with *Sulfolobus* strain Gθ infected with SSV8 and SSV9 at an MOI = 1 (Fig. 7). Two flasks were harvested every 12 hrs from the suite of replicates to determine virus particle counts via ESI/MS. Sulfolobus strain Gθ infected with SSV8 showed typical Gompertzian-like host growth with a concomitant increase in virus particle count (Fig. 7A) was observed as the SSV8 infection continued through 78 hours post-infection (HPI). SSV9-infected Sulfolobus strain Gθ showed a distinct pattern from the Gθ-SSV8 infection. SSV9-infected growth peaked (18 HPI) followed by a rapid drop in cell density (Fig. 7B). The decrease in host density was followed by a spike in SSV9 particle count (32 HPI).

**FIGURE 7.**
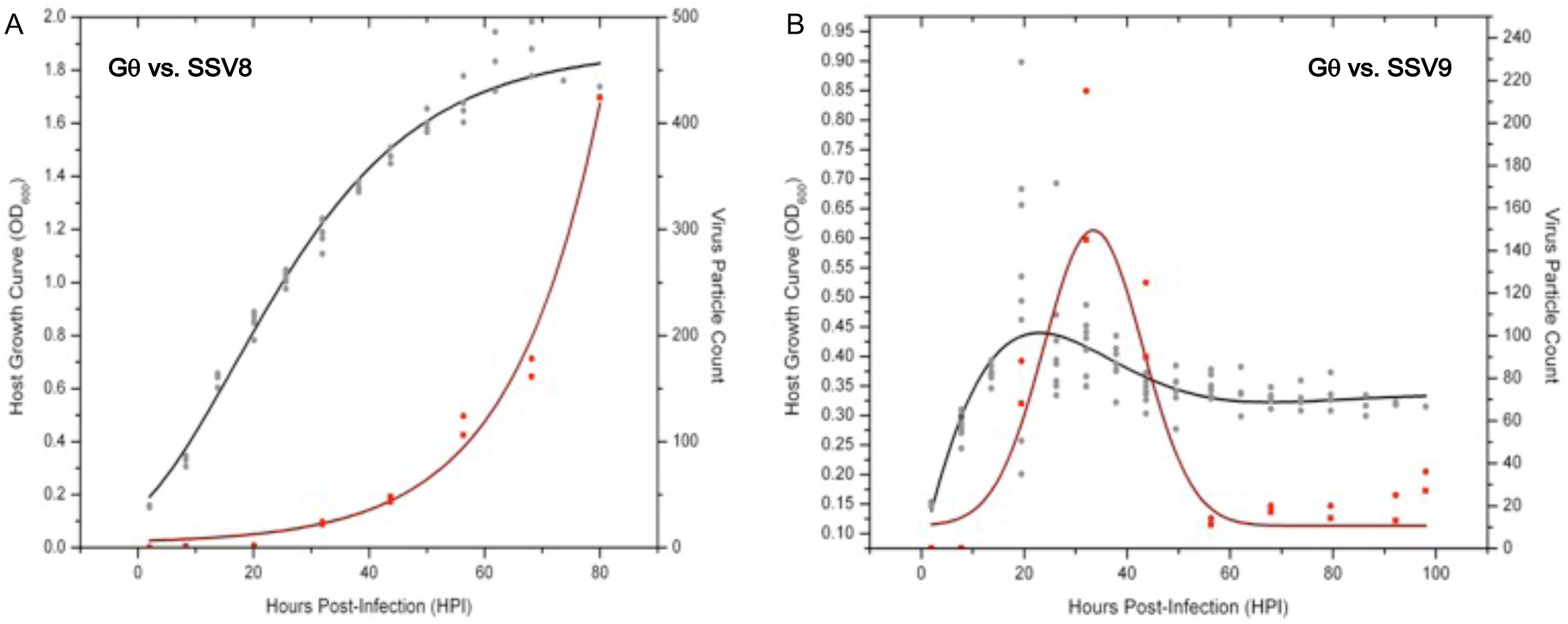
Virus-Host Dynamics: SSV8 versus SSV9 infections in *Sulfolobus* strain Gθ. Using a high multiplicity of infection (MOI=1) to ensure virus particle detection via ESI/MS, strain Gθ was infected with SSV. (A) Sulfolobus strain Gθ at an OD_600_ = 0.15-0.20 is inoculated with SSV8 at MOI=1 resulting in standard Gomperz-like culture growth with a concomitant increase in virion count over time. (B) Sulfolobus strain Gθ at OD_600_≈0.15-0.20 is inoculated with SSV9 at MOI=1 resulting in an average peak density of OD_600_≈0.45 followed by a decrease in cell density and a sudden spike in SSV9 particle count, peaking at 32 HPI. Cell density stabilizes at OD_600_≈0.30-0.35 from 60-90HPI. SSV9 particle count drops to the detectable limits of the ESI/MS (approximately 1×10^6^-2×10^6^ virus particles/mL).

Interestingly, other than the initial peak in host cell density, no noticeable subsequent peaks (dampened or accentuated) were observed in this trial with the higher MOI (*see* Fig. 7B). It was also noted that the growth inhibition did not result in an optical density value of zero. Instead, cell density stabilized at approximately one-third the average peak value. Furthermore, SSV9 titer (via ESI/MS) decreased substantially (*see* Fig. 7B, 60-100HPI). Although the general growth profiles were consistent with prior trials for non-lytic versus lytic release, these latter two features were not expected (*see* Discussion). Together, these data indicate that SSV8 pursues canonical non-lytic virion release expected of SSVs, while SSV9 lyses host strain Gθ. This is also supported qualitatively by the presence of significantly more cell debris in SSV9-infected small-scale infection assays (Fig. S2).

### SSV9 Lytic Replication may be Limited to Allopatric Hosts

To determine if SSV9 lytic replication was dependent upon host strain allopatry, two isolates of Sulfolobales derived from volcanic hot springs in Kamchatka, Russia were also tested as plausible hosts for SSV9. Isolate S147 was derived from the same geothermal area in Geyser Valley as the original host from which SSV9 was isolated. Isolate S150 was derived from the Mutnovsky volcanic region near the south tip of the Kamchatka peninsula. To determine how SSV9 infection would impact growth on a sympatric host (S147) and a *quasi*-sympatric host (S150), spot-on-lawn and liquid culture infection assays were conducted with these two isolates. Spot-on-lawn assays indicate that SSV9 does not form halos on S150 lawns; however, halos readily form on lawns of S147 (Ceballos et al., 2012). Liquid culture-based infection assays substantiate S150 resistance to infection (Fig. 8A) and confirm that S147 is susceptible to SSV9. Post-infection titers for the S150 infection trials did not show any detectable virus; while those for S147 showed virus particle counts on the order of 10^9^ VP/mL indicating productive infection (Fig. S4). The most striking result was that S147 growth during SSV9 challenge resembles the non-lytic archetype infection dynamics observed with other SSVs (Fig. 8B).

**FIGURE 8.**
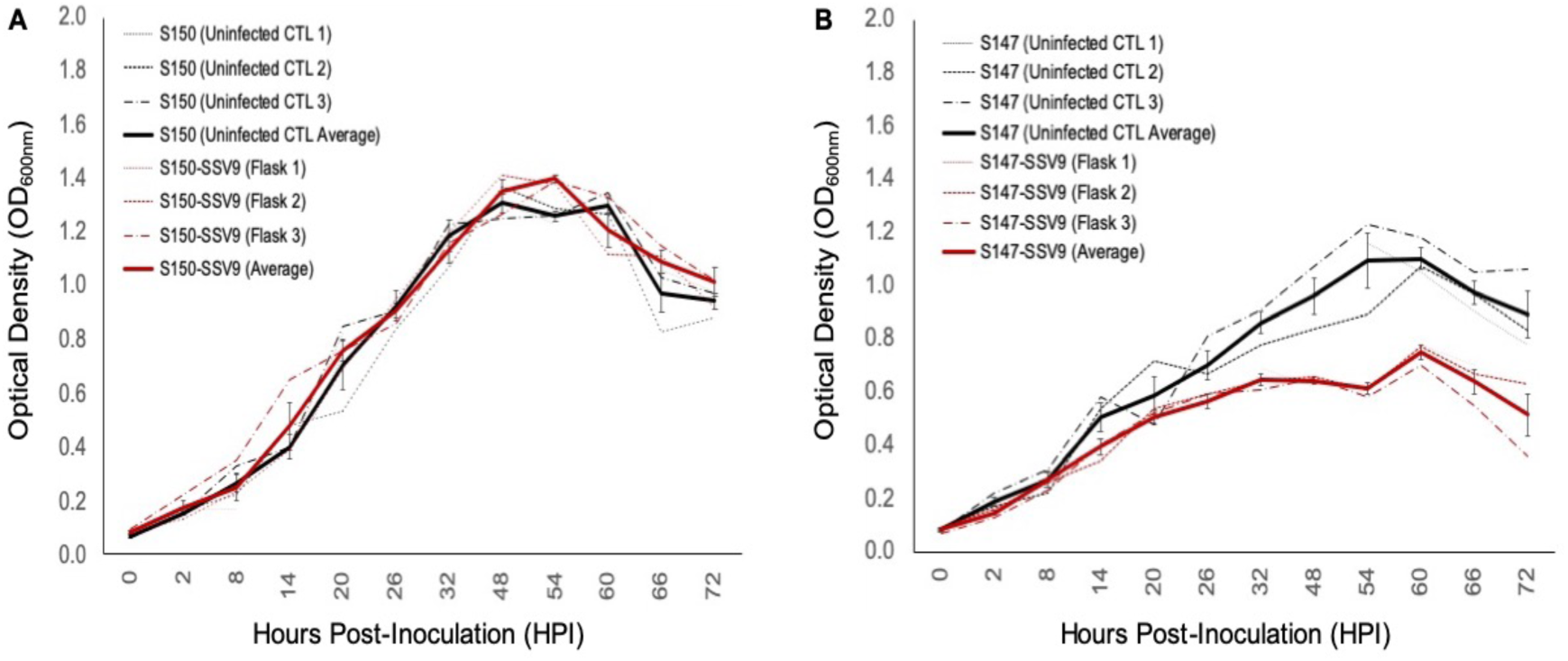
Impacts of SSV9 challenge on Isolates from Geothermal Regions in Kamchatka. Isolates of Sulfolobales from geothermal regions of the Kamchatka peninsula were infected with SSV9. Isolate S147 was derived from the same hot spring region as the host from which SSV9 was derived. S147 is considered a sympatric host. Isolate S150 was also derived from the Kamchatka peninsula but from a geothermal region approximately 256 km to the southwest of Geyser Valley. S150 is considered quasi-sympatric. (A) Strain S150 growth curves for SSV9-infected culture exhibit no significant changes in μmax compared to the uninfected control curves. (B) Strain S147 growth curves for SSV9-infected cultures resemble canonical non-lytic replication.

AUC analysis for S150-SSV9 trials results in no significant growth inhibition (PI < 0.05), while the S147-SSV9 trial shows a significant PI (>0.20) with a convincing Gompertz fit (R^2^ = 0.934), indicating that SSV9 exhibits a non-lytic replication profile on a sympatric host, S147 (*see* Table S3).

## DISCUSSION

Previously published work demonstrated that some Sulfolobales are completely resistant to infection by well-characterized SSVs (i.e., SSV1, SSV2, SSV3, SSV8, SSV9, SSV10) while others are susceptible to subset or all of these SSVs in *spot-on-lawn* “halo” assays (Ceballos et al., 2012). Large plaque-like halos were observed on host lawns challenged with SSV9 (at the same MOI as other SSVs) suggesting that SSV9 is more virulent. However, the large clearings on host lawns generated by SSV9 appeared to be true plaques rather than the turbid halos characteristic of SSV infection. This raised the possibility that SSV9 is lysing host rather than using non-lytic “blebbing” as a means of virion release. Although the halo assay is sufficient for determining whether a given host is susceptible (or not) to a specific SSV, it offers limited information regarding the dynamics of infection. In the present study, liquid culture assays and host growth curve analyses were used to test the hypothesis that: *SSV9 employs a lytic replication strategy rather than the canonical non-lytic replication (i.e*., *virus blebbing) characteristic of other SSVs*.

### SSV infection in liquid culture

Growth curve analysis provides insights into the nature of virus-host infection dynamics that are not resolvable through plate-based halo assays. Using host growth curve analyses, it is not only possible to determine percent inhibition and relative virulence of a different SSVs on a host (*see* Fig. 4A) but it is also possible to gain insights into rates of virus replication/transmission as well as virion release strategy. By measuring titers of inocula at the beginning of an infection trial, establishing equivalent MOI for different virus-host pairings, and taking end-point titers, a quantitative assessment of how many virus particles are present in the culture at the time of inoculation versus how many progeny virions are present at the end-point is possible. However, this method of comparing pre-infection titer and post-infection titers may be confounded by differential stability of distinct SSV strains in the high temperature (76-80°C) and low pH (3.0-3.4) culture environment. Although SSV8 and SSV9 appear to be more resilient in solution, SSV1 does not seem to be as stable (Fig. S3). Thus, viral fecundity may be underestimated by virus count since some virus particles (e.g., SSV1 virions) breakdown during the course of the infection assay. Still, end-point virus count is needed to confirm productive infection. Specifically, if the number of virus particles at end-point exceeds the number of virus particles used for inoculation, then this indicates that the infection was productive. However, if virus count at end-point is equal to or less than virus particle count at inoculation, then it can be argued that no infection occurred. Although it can be argued that a highly unstable virus strain could rapidly breakdown yielding a false negative for infection, TEM (*see* Figs. 1 and 3) and spot-on-lawn halo assays (*see* Fig. 2) are used as secondary methods to validate host susceptibility/SSV strain infectivity.

### Archetypal non-lytic SSV replication and host growth profiles

Despite limitations in determining end-point virus count due to differences in virion stability between the different SSVs, liquid culture assays permit a quantitative assessment of relative virulence (*V*_*R*_) by calculating percent inhibition (PI) of each SSV on a given host strain (*see* Fig. 4B). The archetypal non-lytic replication strategy of SSVs allows quantitative assessment of *V*_*R*_ based on comparisons between *N*_*asymptote*_, μ_max_, and/or AUC in the SSV-infected versus the uninfected control states.

### Atypical infection dynamics of SSV9 on allopatric host growth

Archetypal SSV infection (*see* Figs. 4) induces Gompertz-like host growth profiles (Stacy and Ceballos, 2020), indicative of non-lytic infection, which in turn is consistent with halo, instead of plaque, formation on spot-on-lawn plate assays. However, when SSV9 is used to infect a series of susceptible allopatric hosts, atypical infection dynamics emerge in the host growth curve. Specifically, a deep inhibition, which manifests as “saw-tooth” or “serial spike” profile in the host growth curve, occurs suggesting a different replication process (Fig. 5, *red traces*). Although it may be suggested that SSV9 is simply more virulent on the susceptible host, this argument is refuted by the fact that neither a Gompertz or Logistics growth models fit reasonably to the SSV9-infection host growth data (as indicated by very low R^2^ values).

### Atypical infection dynamics of SSV9 are not MOI dependent

Further support for unique SSV9 infection dynamics rests in the inability to elicit the archetypal non-lytic growth profile by manipulating MOI (*see* Fig. 6B). Even under low MOI, SSV9 continues to induce a cyclic spike profile in the host growth curve. Likewise, varying MOI for SSV8 does not change the archetypal non-lytic growth profile to one resembling SSV9 infection. To determine if a change in the SSV8-infect host growth profile could be forced by a concentrated inoculum, a 20:1 virus concentrate was prepared. Although the culture exhibited a strong depression (*data not shown*), the curve still fit a Gompertz Model with a high coefficient of determination (R^2^ = 0.87) and did not exhibit a cyclic spiking profile. Furthermore, virus particle count at the end of the 20:1 viral concentrate trial was 2-fold greater than what was after the initial inoculum, indicating that the depression in the growth curve was not due to lysis-from-without (Delbrück, 1940) but due to productive infection.

### SSV population dynamics support lytic replication

Larger-scale liquid culture assay trials (i.e., greater number of replicate flasks) allow replicates to be harvested periodically during the course of infection to monitor changes in virus titer in parallel to host growth (Fig. 7). Sampling at shorter intervals (i.e., every 4 hrs), infection with SSV8 (a non-lytic replicator) shows an increase in virus particle count concomitant with host growth. However, SSV9 infection of the same host strain (i.e., Gθ) shows sharp peak in virus particle count ∼12 hrs after a peak and a notable rapid drop in host cell density. This is consistent with lytic virus release or a “burst” event (Fig. 7B). Interestingly, even though the infection assay was carried out to ∼98 HPI, no subsequent “peak-and-crash” of either host (or virus) occurred after the first cycle at 20-32 HPI. Moreover, optical density (a proxy for host cell density) remained steady at OD_600nm_≈ 0.3 from 60-98 HPI. The reason underlying the stability of host cells with no additional SSV9-induced lysing or SSV9 virion production is not clear. There is one report suggesting that SSV9 challenge may induce dormancy in viable hosts (Bautista et al., 2015), which may be the reason for sustained absorbance at OD_600nm_. Indeed, dormant cells would not likely support virus production and may render cells resistant to infection. Another report suggests that group II chaperonin complexes may form highly stable networks of chaperonin complex filaments at the intracellular surface of archaeal cells (Trent et al., 2003). Cell membrane-associated HSP complexes as either filaments or two-dimensional arrays can maintain cell shape even if cells are not viable. Under scanning electron microscopy (SEM), cells from infected culture, which were likely non-viable “ghost cells” due to presence of large holes in the cell membrane, maintained a round, lobed three-dimensional structure that could absorb 600nm light (*data not shown*). Whether the absence of subsequent cycles of host recovery, SSV9 infection, and lytic bursting beyond the initial cycle are due to cells becoming dormant or whether these are “ghost” cells requires further study. Nonetheless, host growth curves for SSV9-infected strains display atypical profiles that do not fit to Gompertzian (or Logistic Growth Models) as is expected for non-lytic virion release but rather represent lytic bursts of virion release. Thus, it is clear from both spot-on-lawn and liquid culture assays that SSV9 is lysing host. Small-scale (10^2^ μl tube) infection assays further support this conclusion. Specifically, infection with SSV9 generates significantly more visible cell debris at the same time point of infection compared to infection with other SSVs (*see* Fig. S2), indicating cell lysis.

### SSV9 lytic replication may depend on host allopatry

Although SSV9 lytic behavior does not appear to be dependent upon MOI and seems to be an inherent property of the SSV9 replication strategy, all susceptible host initially tested were *allopatric* hosts isolated from geothermal hot springs thousands of kilometers away from where SSV9 was isolated. Yet, when two Sulfolobales – one from the same geothermal region from which the original SSV9 was isolated and another from a distant spring (ca. 250 km) within the same area (i.e., Kamchatka, Russia) – were challenged with SSV9, unanticipated dynamics emerged. The *quasi*-sympatric S150 strain showed no susceptibility to SSV9 in spot-on-lawn assays. Moreover, in liquid culture assays with S150, there were no significant differences in *N*_*asymptote*_, μ_max_, or AUC between the SSV9-infected and S150 uninfected controls (Fig. 8A). This is not surprising since other Sulfolobales are reported to be resistant to SSV infection. However, when SSV9 was used to challenge the *sympatric* isolate S147, there were unexpected results. Specifically, it appears that SSV9 does not only infect S147 but that the infection follows canonical non-lytic replication characteristic of other SSVs (Fig. 8B).

Recently, a series of reports have proposed a re-organization in the taxonomy of the family *Sulfolobaceae* to include four distinct genera: *Sulfolobus* (Brock et al., 1972), *Stygiolobus* (Segerer et al., 1991), *Sulphurisphaera* (Kurosawa et al., 1998), and the most recently proposed genus *Saccharolobus* (Sakai and Kurosawa, 2018; Tsubio et al., 2018). Whether SSV infectivity and relative virulence is correlated with these newly defined genera remains to be determined. Our lab has demonstrated that SSVs infect select strains belonging in at least two of these genera. Although this re-organization of the phylogenetic tree for the family *Sulfolobaceae* requires the community to revisit previously published work, the non-lytic growth profile observed in the S147-SSV9 infections raises a question of whether sympatric versus allopatric virus-host evolution determines replication strategy. Since lytic replication typically results in greater virulence (than non-lytic replication), there is also the question of whether a switch in replication strategy over evolutionary timescales (i.e., lytic to non-lytic) is part of a more generalizable pattern of attenuation in the virus-host coevolutionary “arms race” (Van Valen, 1973; Dawkins and Krebs, 1979). Furthermore, the genetic substrates underlying the emergence of distinct replication strategies in SSV systems remain unknown. Employing high throughput sequencing methods (e.g., nanopore sequencing) and advanced analytical techniques may resolve the impacts of specific genetic factors on virus-host dynamics and quantify relative virulence between non-lytic versus lytic phenotypes, respectively (Stacy and Ceballos, 2020), may help to address the aforementioned questions and identify genetic elements that predispose viruses to lytic versus non-lytic replication strategy.

## MATERIALS AND METHODS

### Virus preparation

Glycerol stocks of SSV-infected *Sulfolobus* strains stored at −80°C were partially thawed on ice. For most trials, 100μL of infected cell suspension was added to 30mL of YS media (3.0g/l (NH_4_)_2_SO_4_, 0.7g/l glycine, 0.5g/l K_2_HPO_4_, 2.0g/L sucrose, 1.0g/l Yeast Extract, 200.0μl/l 1% FeSO_4_•7H2O, 244.0μl/l 1% Na_2_B_4_O_7_•10H_2_O, 90.0μl/l 1% MnCl_2_•4H_2_O, 11.0μl/l 1% ZnSO_4_•7H_2_O, 2.5μl/l 1% CuSO_4_•5H_2_O, 1.5μl/l 1% Na_2_MoO_4_•2H_2_O, 1.5μl/l 1% VOSO_4_•5H_2_O, 0.5μl/l 1% CoCl_2_•6H_2_O, 0.5μl/l 1% NiSO_4_•6H_2_O, 1.0μl/l 1.0M MgCl_2_/0.3M Ca(NO_3_)_2_ solution; pH adjusted to 3.2 with H_2_SO_4_) in a 125mL Erlenmeyer flask. Alternatively, T media was used in select experiments using the same solution but with 3.0g/l tryptone substituted for the yeast and sucrose components. Flasks were partially sealed to avoid evaporation and incubated at 76-78°C/70-90 RPM. Once liquid culture reached an optical density (OD_600_) between 0.4 and 0.6, 3.0mL of cell suspension was used to inoculate 600mL of pre-heated fresh media in a 1.0L baffled flask. These cultures were incubated at 78°C/70 RPM until reaching an OD_600_ = 0.6-0.8 to maximize virus yields. All 600mL of the culture was centrifuged for 20 min at 6,000 RPM (Sorvall RT Legend Centrifuge, Fiberlite 4×8000ml fixed-angle rotor with 250ml inserts; ThermoFisher, Pittsburgh, PA) to pellet the cell mass while leaving virus in suspension. The upernatant was decanted and filtered through a 0.45μm vacuum PES filter system. The filtrate was concentrated using 10kDa Centricon Plus-70 spin concentration tubes (Millipore Corp., Billerica, MA, USA) to produce ∼3.0mL concentrated SSV suspension. TEM was used to confirm the presence of virions. Virus particle count was measured using electrospray ionization/mass spectrometry or serial dilution plaque-like assays (see below). Virus was stored at 4°C and used in *spot-on-lawn* “halo” assays and liquid culture infection assays within 2-3 weeks of harvesting. Dilutions of viral stocks were used as inocula.

### Host cell preparation

Glycerol stocks of uninfected *Sulfolobus* strains stored at −80°C were used to establish 30mL cultures in YS (or T or TS) media, as described above. Uninfected *Sulfolobus* cultures were either used to prepare host lawns for halo assays (Ceballos et al., 2012) or used in liquid culture infection assays (as described below).

### Transmission electron microscopy (TEM) of viruses and cells

To verify the presence of SSVs in samples, ∼5μL of viral suspension was spotted onto a formvar-coated copper grid and incubated for 10 min in a humidity chamber. The sample was rinsed off with distilled water and negatively stained with a 1% solution of uranyl acetate for 30 seconds. The stain was wicked and the sample was air dried. Grids were imaged with a Hitachi H-7100 TEM at 75kV. Images were captured at 60,000-200,000X magnification. TEM images for cells were acquired as described in Brumfield et al. (2009). Cells were fixed in glutaraldehyde (3% v/v), centrifuged, and re-suspended in a small volume of agar (2% v/v). After solidification, the resulting agar was cut into small pieces and fixed overnight with glutaraldehyde (3% v/v) in 0.05M potassium sodium phosphate buffer (PSPB) and pH=7.2. Agar pieces were rinsed twice for 10 min each with PSPB. Agar pieces were post-fixed with osmium tetroxide (2% v/v) at RT for 4h. Samples were dehydrated via an ethanol rinse series (50%-100% v/v) and then washed with a transitional solvent, propylene oxide. Spurr’s resin (Spurr, 1969) was used to infiltrate dehydrated cell mass and pieces were baked overnight at 70°C. Thin sections (60-90 nm) were cut and stained with uranyl acetate and lead citrate. Imaging was done with a LEO 912AB TEM.

### Electrospray ionization/mass spectrometry (ESI/MS)

Virus suspension preparations were processed in an Integrated Viral Detection System (BVS, Inc.; Missoula, MT) per protocols in Wick et al. (1999). In brief, a mixture consisting of 100μL viral suspension and 900μL ammonium acetate buffer solution is aerosolized in a Electrospray Aerosol Generator (Model 3480, TSI, Inc., Minnesota). A Differential Mobility Analyzer (Model 3081, TSI, Inc., Minnesota) separates particles by their electrical mobility, which is influenced by particulate mass-to-charge ratio or “size”. These particles flow in tandem with a saturated butanol fluid. The particles initiate butanol condensation and the stream is cooled enabling butanol-condensed particles to be optically counted in a Condensate Particle Counter (Model 3772, TSI, Inc., Minnesota). System software displays results in terms of particle count per size category with a standard range of 2-280nm. This specialized ESI/MS is designed to detect intact virus particles (Wick et al., 2006) and can measure relative virus particle count between two or more samples. In these infection assays, ESI/MS spectra were evaluated by using either peak values or the area under the curve to compare SSV particle production between two cultures.

### Note

Time-of-flight estimation based on mass:charge is used to calculate ESI/MS “size”. ESI/MS “size” will differ from physical size as measured by other methods such as transmission electron microscopy. In addition, mass:charge size versus physical size will differ due to the assumption of virus particle sphericality in time-of-flight derivations. SSVs have been shown under electron microscopy to be fusiform or “spindle-shaped” particles of approximately 60 × 90 nm (and not spherical), thus, a discrepancy is expected. In some cases, SSVs exhibit dual peaks or shouldered peaks under ESI/MS. Such spectral features may be due to different conformations of the same SSV or, potentially, due to the presence of pleomorphic SSVs. Under ESI/MS, SSVs exhibit characteristic peaks between 46.1 and 61.5 nm for purified virus suspensions with major peaks typically appearing at 46.1, 47.8, and 49.6 within the ±4nm system tolerance.

### Spot-on-lawn halo assays

Halo assays were performed as in Stedman et al. (2003). Cultures were grown as described above. At an OD_600_= 0.4-0.6, 500μL of cell suspension was added to 4.5mL of a 78°C mixture of equal parts 1.0% w/v Gelrite^®^ (Sigma-Aldrich, St. Louis, MO, USA) and two-fold concentrated YS medium. The 5mL mixture was spread on pre-warmed Gelrite^®^ plates (1% w/v in medium) and allowed to solidify for 15 minutes at room temperature followed by a 20 min incubation at 78°C. 1.0μL of each viral suspension was spotted onto the prepared plate in labeled areas. Depending upon host strain, plates were incubated for 3-9 days at 78°C. Successful infection was scored by the formation of a visible *halo* of growth inhibition on the host lawn. Triton X-100 (0.05% v/v) was used as a positive control and sterile water was used as a negative control.

### Liquid culture infection assays

As described above, 3.0mL of uninfected *Sulfolobus* culture was diluted 1:100 in pre-heated media in a 1.0L baffled flask. Cultures were grown to an OD_600_≈ 0.15. Then, 100μL of standardized viral suspension was added to the flask. *Sulfolobus* cultures were incubated with shaking (78°C/70RPM) for various time intervals. Virus was then harvested as detailed above and ESI was used for virus particle counts. Growth curves from SSV-*Sulfolobus* cultures were fit to: Logistic and modified Logistic *(eqs. 1,2)*; Gompertz and modified Gompertz *(eqs. 3, 4)*; and, other growth models (Gompertz, 1825; Laird, 1964; Lopez et al., 2004, Zwietering et al., 1990; Sprouffske & Wagner, 2016):

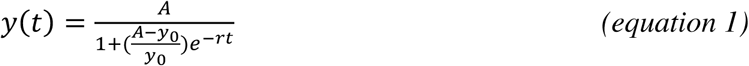

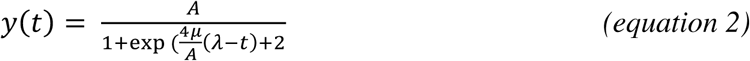

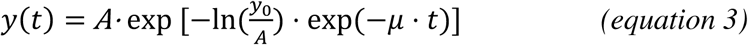

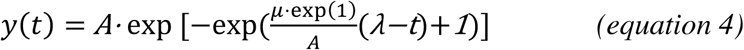

The *exp* represents an exponential function such that exp(x)=e^x^, where e is Euler’s number. *A* represents the amplitude or peak growth value in the given environment, which corresponds to stationary phase and the maximum carrying capacity; *r* is the intrinsic growth rate coefficient; *y*_0_ is the initial population size; μ is the maximum slope of the growth curve, and *t* is time. For equations in which the maximum specific growth rate, μ or μ_max_, was not a fit parameter; it was calculated by taking the derivative of *y*(*t*) evaluated at a software-derived point of max growth using parameter estimates for A, *r*, and *y*_0_.

Parameter estimates were derived in R 3.6.1 (R Core Team; R Foundation for Statistical Computing, Vienna, Austria) using curve fitting functions of easynls, growthcurver, and grofit packages (Arnhold, 2017; Sprouffske & Wagner, 2016; Kahm *et al*., 2010). The best parameter values for each function were found via Levenberg-Marquardt non-linear least-squares algorithm (Moré, 1978). The levels of fit between software packages varied and provided varied slightly different parameter values for each dataset. Relative area under the curve values; however, remained consistent across assessments within the same fitted models (i.e. Gompertz and Gompertz with software “A”) and with direct calculation of Riemann sums of integral of curves. The area under the optical density-based growth curves was calculated via two methods for each data set. The fitted models were integrated with from time *t* = 0 to the time-point corresponding to the end of the exponential growth phase of the uninfected host-control’s growth curve. This process was utilized for each data set was able to be successfully fit to equations 1-4 above, as well as other growth models not shown here. The second method for calculating AUC values was through Riemann sums, also known as the trapezoidal approximation method (Hasenbrink *et al*., 2005) Equation 5 shows the formula utilized for calculating the AUC with this method.

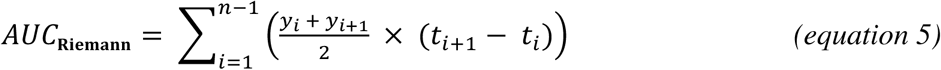

Area under the curve values for virally challenged hosts were then standardized to the AUC value for the averaged uninfected host control growth curve in order to calculate the percent inhibition (PI) for the individual growth curves (*see* Tables S1, S2, and S3). The equation used to calculate the percent inhibition of growth is shown in equation 6 below, from (Rajnovic *et al*., 2019) which is reformatted in equation 7. For a full discussion of this method (*see* Stacy and Ceballos, 2020).

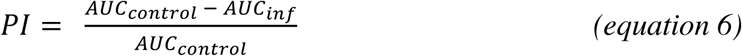

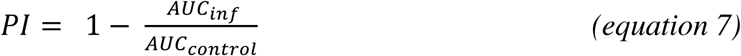

The left and right bounds of the integration process are key parameters affecting the area under the curve metric. For these viral growth curves, the point of first difference is at time t = 0 when virus is added. The upper bound of the growth curve was originally chosen to be the intercept of the line produced by the maximum growth rate and the fitted asymptotic value. This method was opted against because it does not include the transition period between the exponential growth phase and the stationary phase of growth, an area of the growth curve which we considered to be worthy of inclusion. Datapoints at and beyond stationary phase were excluded from AUC analysis because they disproportionately value the asymptotic (A) parameter, which undermines the robustness of the AUC value as a unifying metric of growth curve changes.

Virus replication data, for SSV8, was fit to an exponential growth model both based on peak values and AUC calculations (with similar patterns emerging with either approach):

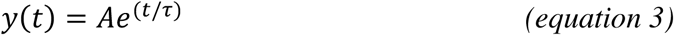

Parameter A is the initial population prior to infection and 1/τ is the growth rate constant. For SSV9-infected cultures, average OD-based growth was fit to a sinusoidal function:

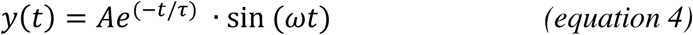

Term Ae^-t/τ^ represents the exponential decay of the oscillation amplitude whereby A is the driving amplitude and 1/τ is a decay rate constant. In the second term, ω is the frequency variable. The average growth curve of SSV9-infected cultures was fit to a damped sinusoidal function simply because this function best described the cyclical bursts in growth that became increasing smaller in terms of amplitude and which were each followed by rapid declines in host population (as determined by optical density readings). However, for SSV9 replication, a Gaussian function was used to fit the dataset.

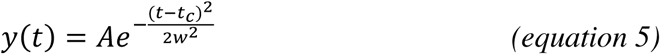

Parameter A represents the amplitude of the peak function; t_c_ is the time at which the center of the peak occurs; and, w represents the width of the peak function at one-half the peak amplitude. The “goodness-of-fit” of these data sets (assessed by adjusted r^2^ values) to the sinusoidal and Gaussian, respectively, prompted their use. For the sinusoidal fit, the adjusted r^2^ value was 0.90 (*see* Fig. 7B). For the Gaussian fit to the SSV9 virus replication data, adjusted r^2^ = 0.94. Note that r^2^ values typically were between 0.90-0.99, except for fits to the SSV9-infected host growth curves, which failed or had r^2^ < 0.25.

### qPCR for determination of SSV9 genome copy number

SYBR green-based qPCR was used to quantify the number of SSV9 genomes for inocula and end-point infection. Two PCR primers were employed: SSV9F (5’-GTGAAGCGACCAACATAGGTGCAA-3’) and SSV9R (5’-GTTGCGTTTGTACCGGTTACGCTA-3’) - targeting the single-copy gene *vp1*, which encodes for a major SSV structural capsid protein (Bautista, 2015). Standard curves were generated with 10-fold serial dilutions (108 to 100 copies) of the *vp1* gene fragment (138-bp) cloned into a TOPO TA pCR2.1 plasmid (Invitrogen). Copy number in each standard was calculated using formula:

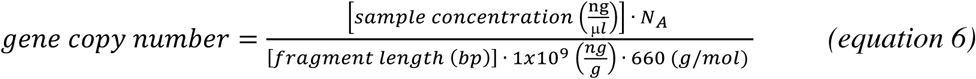

where, 660 g/mol is the molecular weight of one base pair, *N*_*A*_ is Avogadro’s number, and 1×109 is used to convert units to nanograms. Each qPCR reaction was performed on a Rotor-Gene Q (Qiagen, Inc.; Maryland, USA) and consisted of 1X Green/ROX qPCR Master Mix (Fermentas, Inc.; Ontario, Canada), 3 pmol of each primer, 1 uL of DNA template, and nuclease-free water up to a final volume of 20 uL. Three technical replicates were performed of each standard and samples. The qPCR cycle parameters were as follows: 98°C for 2 minutes, followed by 40 cycles of 98°C for 5 s and 20s at 60°C. A melt curve analysis was performed after each run from 65 to 95°C in 0.5°C increments at 2 s intervals, to ensure specific amplification of a single target, and no primer dimer formation. The qPCR amplicons were resolved in 2% agarose gels to assess amplicon sizes for target-specific amplification. Reaction mixtures with sterile water (no-template DNA) or pCR2.1 DNA without insert served as a negative controls to control for false positives. Assays showed amplification efficiencies of 100% ± 10% (i.e., with a slope between −3.6 and −3.1), consistency across replicate reactions, and linear standard curves (*R*2 > 0.970). The absolute quantification method was used to calculate the number of viral genomes per mL of culture.

## AUTHOR CONTRIBUTIONS

RMC and KS conceptualized the study. RMC performed liquid culture and plate assays. RMC did virus titering using ESI/MS with instrumentation at BVS, Inc. (Hamilton, MT). CD performed plate assays and conducted the SSV thermal stability and adsorption studies. CD also did virus titering using serial dilution plaque assays. CLS and RMC fit data to mathematical models and determined N_asymptote_, μ_max_, and AUC for all liquid culture assays. RMC and CD produced TEM and SEM images. EP did qPCR-based virus titers.

## ACKNOWLEDGEMENTS

The authors thank: UA students Kyle Hanson and Blythe Bunkers for assisting with infection assays; David Wick (BVS, Inc.) for technical support with the ESI/MS titering; and, Dr. Jonathan Trent and Dr. Mark Young for providing several strains of Sulfolobales derived from different geothermal regions worldwide for comparative analyses.

## SUPPLEMENTARY MATERIAL

**FIGURE S1.**
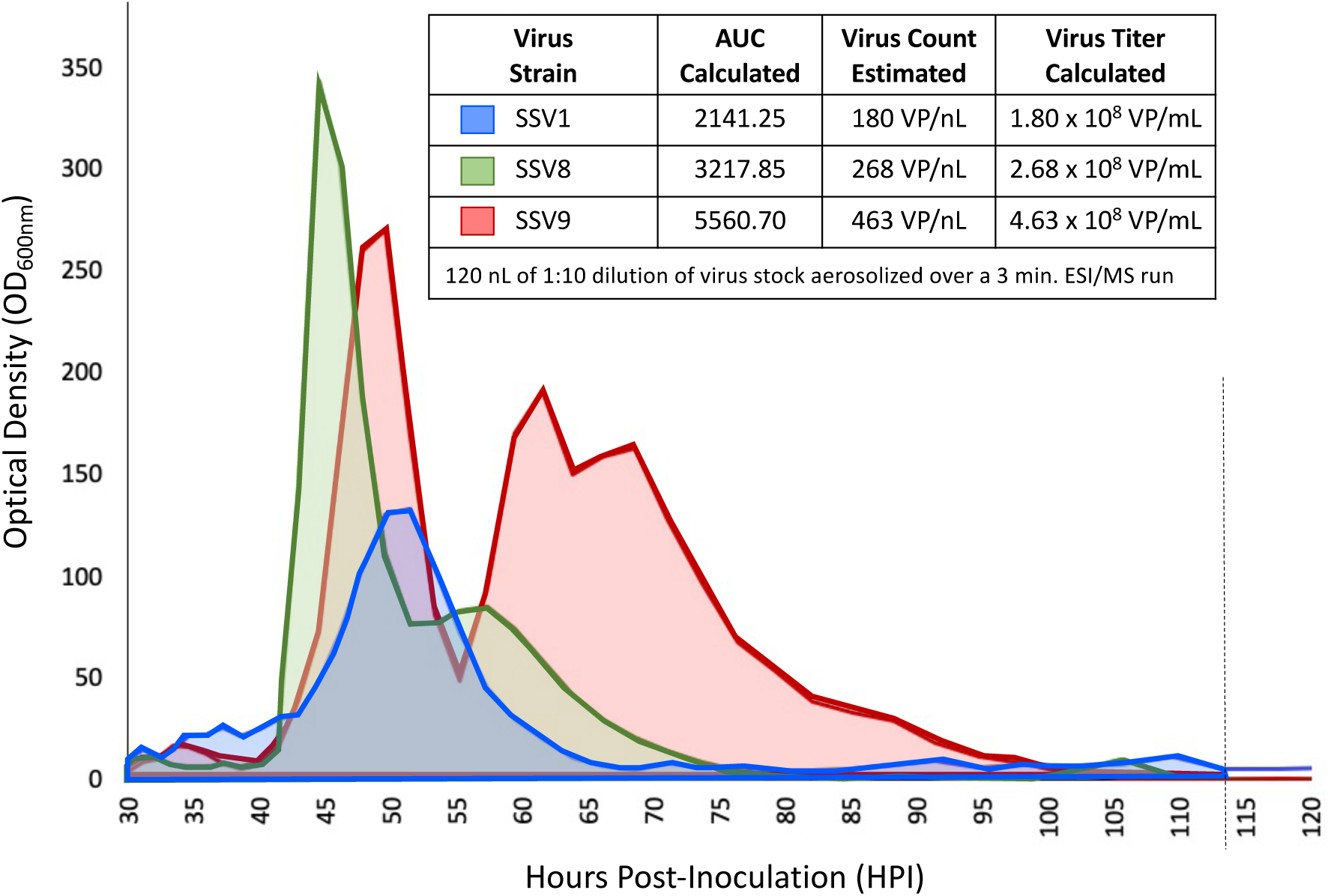
SSV Titers via Electrospray Ionization Mass Spectrometry (ESI/MS). Concentrated viral suspension is diluted 1:10 and mixed with ionization solution (*see* Methods section). SSV particle count in 120nL of suspension is monitored by a particle detector for 3 min. The area-under-the-curve (AUC) provides an estimate of titers. Each SSV has a characteristic spectrum. SSV1 (blue) is generally a single broad peak. SSV8 (green) exhibits a large single peak with a shoulder. SSV9 (red) typically presents as a double-peak. Since ESI/MS measures are based on mass:charge ratios of spherical particles and SSVs are fusiform (quasi-elliptical) shoulders and double peaks may be due to different orientations during between aerosolization and triggering the detector. Alternatively, morphologically different populations may be present, including a sub-population of defective virus-like particles such as those that have lost their hydrophobic tails, which generate a shoulder or second peak.

**FIGURE S2.**
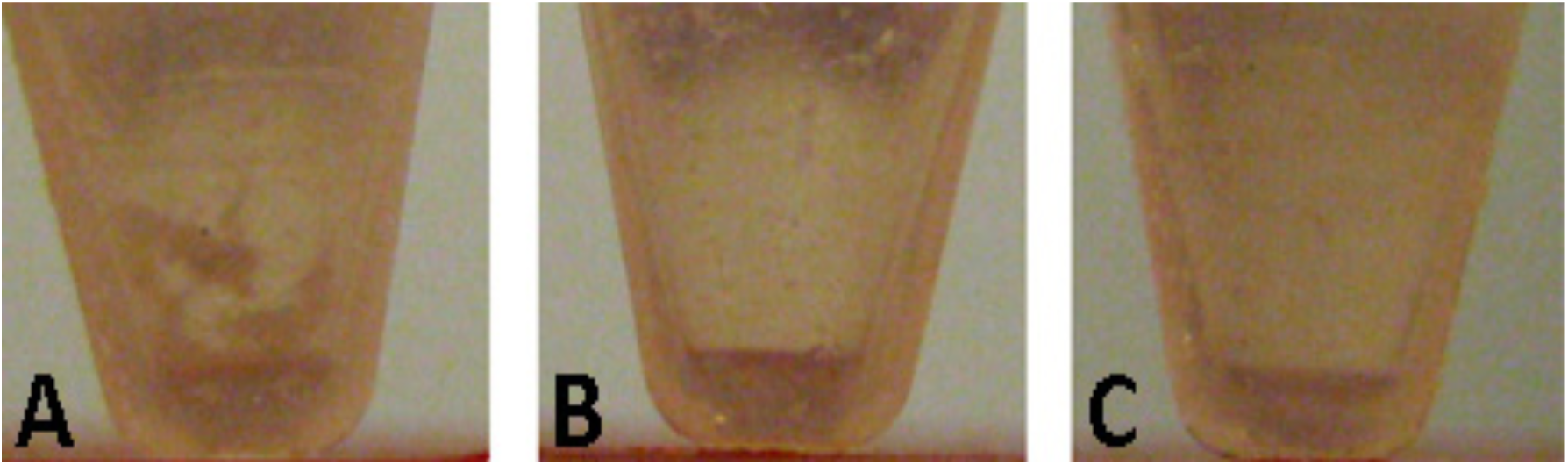
Debris in SSV9-infected Small-Scale Cultures. (A) SSV9-infected *Sulfolobus* strain Gθ shows significantly more cell debris at 22HPI than (B) SSV-infected Gθ at 22 HPI and (C) uninfected Gθ uninfected control. Both infections were conducted at an MOI=3 where virus titer was determined by plate-based serial dilution plaque-like “halo” assays (units = halo-forming units/mL or hfu/mL). Cell debris is another line of evidence suggesting SSV9-induced host cell lysis. *S.solfataricus* Gθ infected with SSV1. **C)** *S.solfataricus* Gθ without the addition dark region at the bottom of all three tubes is a reflection. Contrast was automatically Photoshop CS4 and all three tubes are from the same photograph. Tubes have centrifuged or otherwise treated.

**FIGURE S3.**
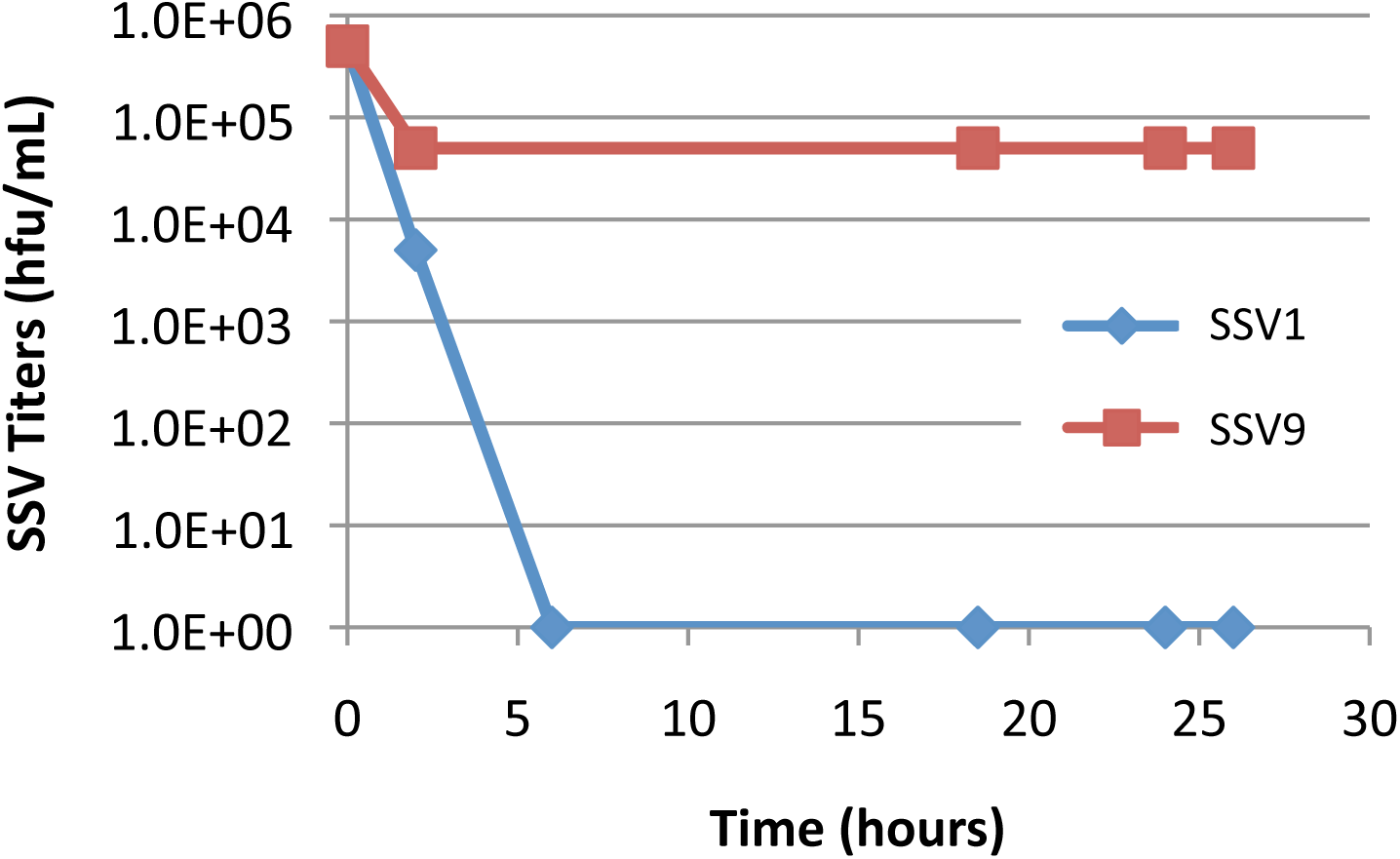
SSV Viability After Exposure to Culture Environment. At 75°C (pH 3.2) SSV9 (red) retains viability for up to 25 hours; while titers of SSV1 (blue) decrease rapidly after 6 hr in the same high temperature acid conditions. Plate-based serial dilution plaque-like “halo” assays (hfu/mL) were used to determine titers for each time point for SSV9 and SSV1. SSV9 virus particles (red) have greater stability in the high temperature and acid environment. determined via halo assay. Both SSV-1 (blue diamonds) and SSV-(red squares) exhibited initial drops in titer, though the SSV1 sample had no detectable virus (<500 hfu/mL) after 6 hours, while SSVK-1 retained infectivity after 26 hours of incubation. Adsorption timing: Accurate measurement of virus production by newly infected cells requires washing away excess viruses used in the initial infection, following their adsorption to cells. To ensure that cells were not washed prior to infection taking place, adsorption time was determined indirectly by loss of virus titer in the presence of Sulfolobus solfataricus Gθ cells at an MOI 0.01, (100 fold excess cells). Because virus titers of concentrated SSV1 and SSVK-1 stocks decreased significantly at 75°C in the absence of cells (Figure 2), this assay was performed 23°C to analyze the loss in virus titer due to the presence of cells independent from the loss

**FIGURE S4.**
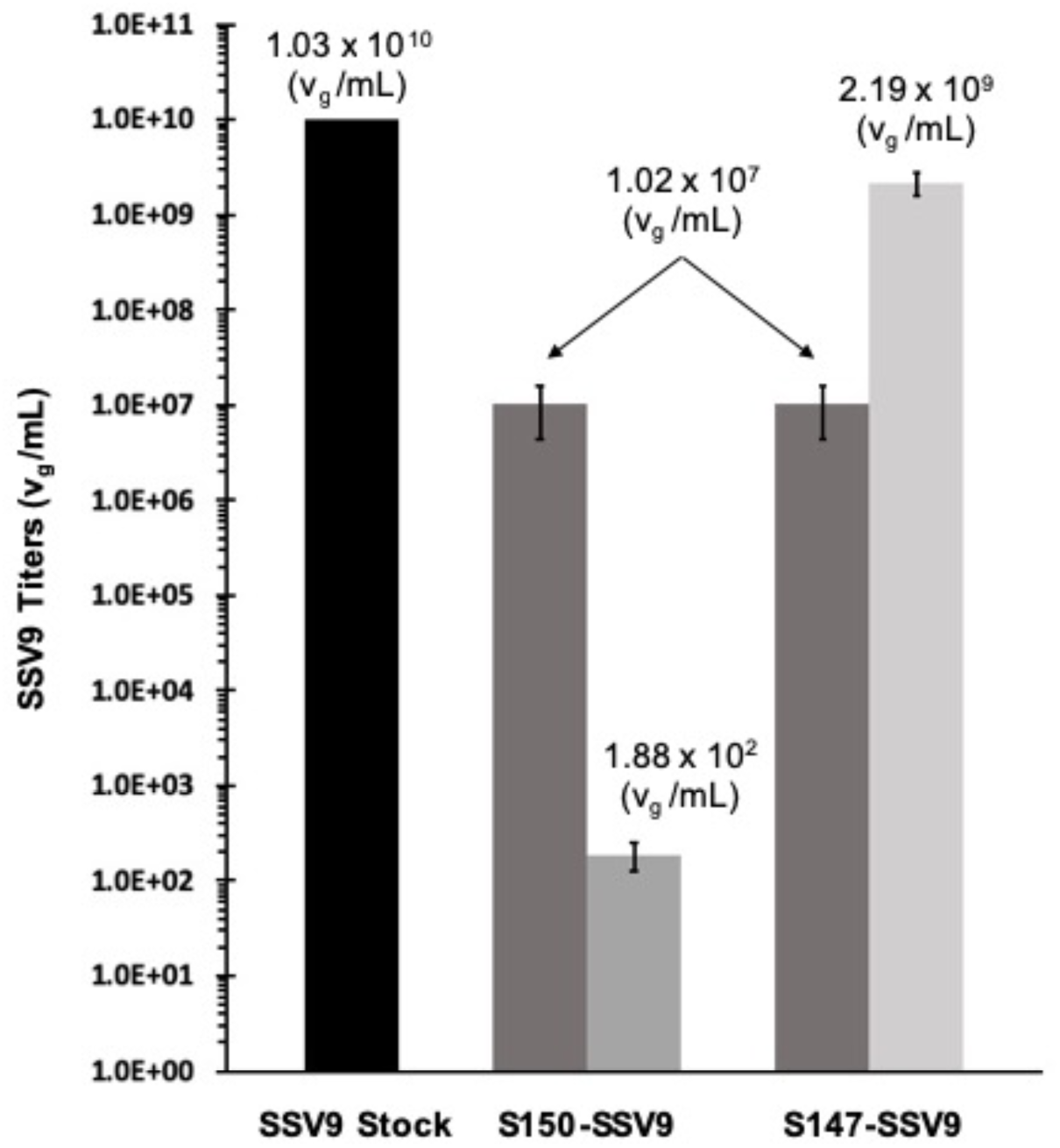
SSV9 Genome Abundance via Quantitative Polymerase Chain Reaction (qPCR). Viral genome counts were measured using qPCR For S150-SSV9 and S147-SSV9 challenges. Inoculation of S150 and S147 cell suspensions with SSV9 at an MOI = 0.1 (dark grey bars) from a SSV stock titered at 1.03×10^10^ v_g_/mL (black bar) was followed by incubation for 72 hours at 78°C (90 RPM shaking). At end-point 72 HPI, titers were determined by qPCR from culture supernatant. The presence of SSV9 genomes per mL in S150 culture dropped five orders of magnitude to within the lowest detectable limits of the system (medium grey bar), which indicates no productive infection. SSV9 genomes in S147 culture increased by two orders of magnitude (light grey bar), indicating that S147 is permissive to SSV9 replication.

**Table S1.**
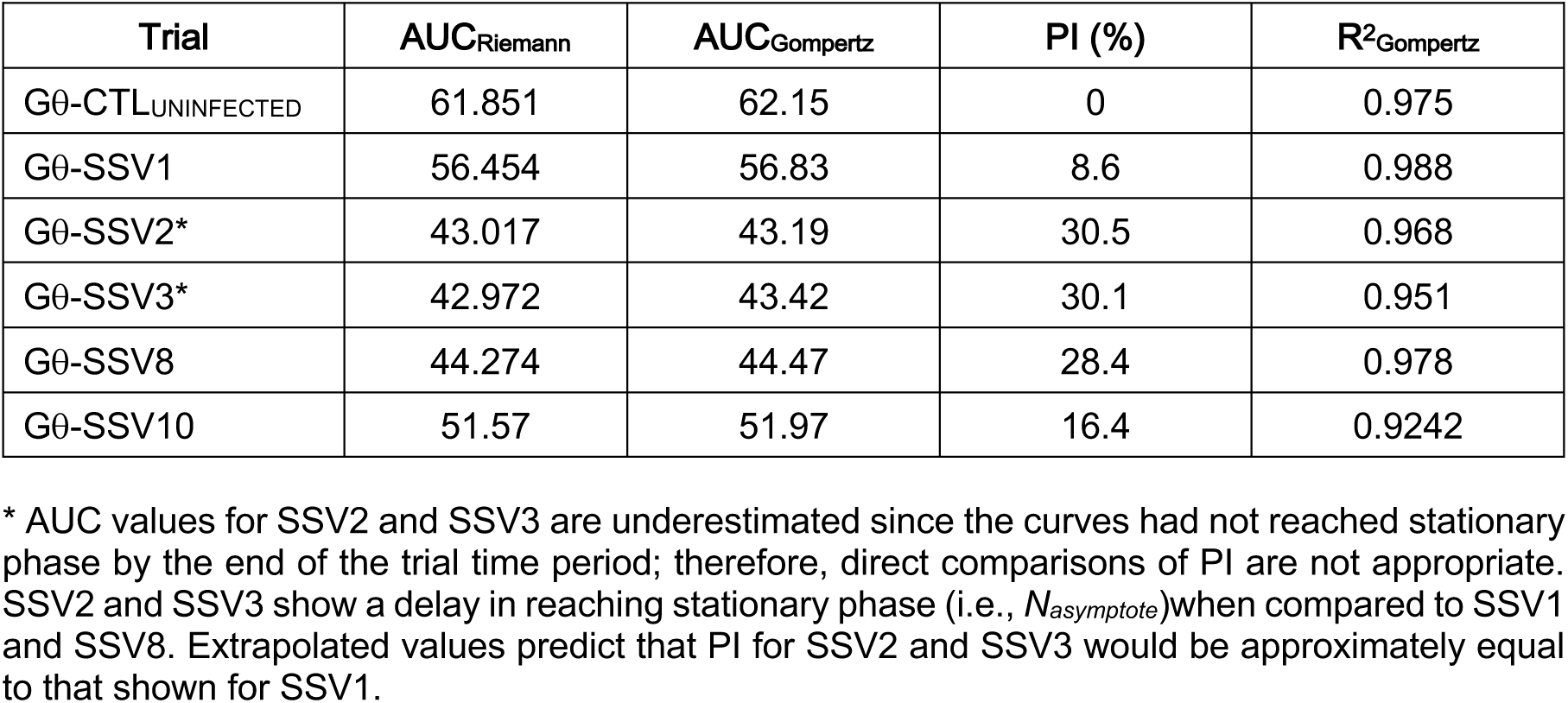
Percent Inhibition Calculations for SSV-Gθ Infection: Figure 4.

**Table S2.** Percent Inhibition for SSV Infections on Allopatric Hosts: Figure 5.

|  |  |  |  |  |
| --- | --- | --- | --- | --- |
| **Trial** | **AUC** | **PI (%)** | **R<sup>2</sup><sub>Gompertz</sub>** | **SE<sub>AUC</sub> (±)** |
| S200-CTL<sub>AVG</sub> | 113.4 | 0 | 0.978 | 0.792 |
| S200-SSV1<sub>AVG</sub> | 107.5 | 5.3 | 0.986 | 4.104 |
| S200-SSV8<sub>AVG</sub> | 75.0 | 33.9 | 0.965 | 0.691 |
| S200-SSV9<sub>AVG</sub> | 31.4 | 72.4 | **0.301** | 1.332 |
| S444-CTL<sub>AVG</sub> | 70.3 | 0 | 0.976 | 1.778 |
| S444-SSV1<sub>AVG</sub> | 59.6 | 15.1 | 0.980 | 1.070 |
| S444-SSV8<sub>AVG</sub> | 51.8 | 26.3 | 0.970 | 0.357 |
| S444-SSV9<sub>AVG</sub> | 24.2 | 65.6 | **0.479** | 1.172 |
| S437-CTL<sub>AVG</sub> | 102.6 | 0 | 0.962 | 0.221 |
| S437-SSV1<sub>AVG</sub> | 99.0 | 3.5 | 0.981 | 1.603 |
| S437-SSV8<sub>AVG</sub> | 69.9 | 31.8 | 0.969 | 6.272 |
| S437-SSV9<sub>AVG</sub> | 31.6 | 69.2 | **0.531** | 4.728 |
| Coefficients of Determination (R<sup>2</sup>) less than 0.8 (in **bold**) are considered “failed” fits to the Gompertz model.<br>For strain descriptions and references see main text. |  |  |  |  |

**Table S3.** Percent Inhibition for SSV9 Challenges on Sympatric Sulfolobales: Figure 8.

|  |  |  |  |  |
| --- | --- | --- | --- | --- |
| **Trial** | **AUC** | **PI (%)** | **R<sup>2</sup><sub>Gompertz</sub>** | **SE<sub>AUC</sub> (±)** |
| S150-CTL<sub>1</sub> | 54.1 | -1.5 | 0.935 |  |
| S150-CTL<sub>2</sub> | 51.5 | 3.5 | 0.857 |  |
| S150-CTL<sub>3</sub> | 54.4 | -1.9 | 0.924 |  |
| S150-CTL<sub>AVG</sub> | 53.3 | 0 | 0.920 | 0.92 |
| S150-SSV9<sub>1</sub> | 53.3 | 0.0 | 0.931 |  |
| S150-SSV9<sub>2</sub> | 54.5 | -2.2 | 0.913 |  |
| S150-SSV9<sub>3</sub> | 55.8 | -4.7 | 0.941 |  |
| S150-SSV9<sub>AVG</sub> | 54.5 | -2.3 | 0.942 | 0.72 |
| S147-CTL<sub>1</sub> | 42.7 | 1.0 | 0.917 |  |
| S147-CTL<sub>2</sub> | 39.9 | 7.3 | 0.946 |  |
| S147-CTL<sub>3</sub> | 46.7 | -8.3 | 0.955 |  |
| S147-CTL<sub>AVG</sub> | 43.1 | 0 | 0.967 | 1.97 |
| S147-SSV9<sub>1</sub> | 31.8 | **26.3** | 0.912 |  |
| S147-SSV9<sub>2</sub> | 31.4 | **27.2** | 0.960 |  |
| S147-SSV9<sub>3</sub> | 30.4 | **29.6** | 0.843 |  |
| S147-SSV9<sub>AVG</sub> | 31.2 | **27.7** | 0.934 | 0.41 |
| For strain descriptions and references see main text. |  |  |  |  |

